# Transcriptome of Peripheral Blood Mononuclear Cells Reveals Suppressed MAPK/AP-1 Pathways During *Ascaris-Salmonella* Coinfection in Pigs

**DOI:** 10.64898/2026.01.24.701481

**Authors:** Zaneta D. Musimbi, Rima Hayani, Robert M. Mugo, Larissa Oser, Josephine Schlosser-Brandenburg, Sebastian Rausch, Susanne Hartmann, Ankur Midha

## Abstract

*Ascaris* and *Salmonella* are prevalent pathogens in pigs, and their coinfection could pose significant veterinary and public health concern. While *Salmonella* typically elicits strong monocyte-driven inflammation, we previously showed that *A. suum* coinfection impairs monocyte responses and increases bacterial burden. Building on these prior observations, we investigated the transcriptional basis of helminth-induced immune modulation using peripheral blood mononuclear cells from experimentally infected pigs. Bulk RNASeq analysis revealed 126 differentially expressed genes in coinfected pigs relative to *Salmonella* single-infected pigs, including downregulation of genes associated with chemotactic function (*CCL3L1*, *CCL8*, *CXCL14*) linked to monocyte recruitment and macrophage-mediated antimicrobial function. To uncover underlying cellular signaling mechanisms, we applied co-expression network analysis, identifying two modules of interest: one enriched for inflammatory signaling pathways (TNF, IL-17, MAPK), and the other associated with phagosome and lysosome function. Notably, coinfection resulted in selective repression of key genes in the inflammation-related module, including MAPK modulators (*DUSP1, DUSP6*), AP-1 components (*FOS, NR4A1, MAFF*), and monocyte activation genes (*TNFSF9, CD163*), pointing to a potential coordinated shutdown of monocyte inflammatory signaling. These findings reveal that an active *Ascaris* infection interferes with host immunity against a subsequent bacterial infection by disrupting AP-1/MAPK-dependent transcriptional networks, providing mechanistic insight into helminth-mediated immune modulation.

## Introduction

Nontyphoidal *Salmonella* infects approximately 150 million people worldwide with 60,000 deaths reported each year ^1, 2^. While pigs are generally considered as asymptomatic carriers of nontyphoidal *Salmonella,* they are among the major causative agents of human foodborne salmonellosis ^3–5^. Equally relevant to global human public health is *Ascaris lumbricoides*, a soil-transmitted helminth highly prevalent in low- and middle-income countries ^6, 7^, associated with malnutrition, anemia and impaired cognitive development in children ^8–11^. The porcine roundworm, *Ascaris suum,* genetically closely related to *A. lumbricoides* ^12–14^ and is associated with significant economic losses in pig farming with zoonotic potential ^15^.

Nontyphoidal salmonellosis is mainly an enteric disease that triggers a Th1/Th17 immune response in pigs ^16, 17^. Mouse model studies have highlighted the critical role monocytes play in the innate and adaptive response against salmonellosis ^18^. The early phase of infection induces an inflammatory response, and chemokine-mediated recruitment of neutrophils and inflammatory monocytes derived from the bone marrow to the gut-draining lymphoid tissues ^18, 19^. These monocytes differentiate into macrophages, crucial for phagocytosis and controlling bacterial replication ^19, 20^. Additionally, the recruited monocytes and monocyte-derived macrophages secrete proinflammatory cytokines like TNF-α and IL-1β, which enhance bactericidal activity ^19, 21^. While classically activated macrophages induce antibacterial proinflammatory cytokines, alternatively activated macrophages provide a conducive environment for *Salmonella* replication ^21, 22^. In contrast, ascariasis is associated with a hepato-tracheal migration that induces localized Th2 immune responses ^23–25^. In a murine model, we and others have shown that the larval migratory phase of ascariasis induced neutrophilia followed by eosinophilia and eventually recruitment of monocytes that differentiate into alternatively activated macrophages in the local sites of infection; the liver and the lungs ^26, 27^.

Our group recently investigated the impact of larval ascariasis on anti*-Salmonella* responses in experimentally coinfected pigs ^24^. We found that coinfected pigs exhibited higher *Salmonella* burden in the jejunal mesenteric lymph nodes compared to *Salmonella* single-infected pigs. Among others, this was accompanied by reduced monocyte frequencies in peripheral blood mononuclear cells (PBMCs) and a diminished monocyte TNF-α response following IL-12 and IL-18 *ex vivo* stimulation as well as higher susceptibility of macrophages to *Salmonella* infection *ex vivo* ^24^. Additionally, in a different study, we showed that exposure to *A. suum* excretory/secretory (ES) products *in vitro* reduces monocyte TNF-α production ^28^, supporting a helminth-driven immunomodulatory effect.

These findings suggest that *A. suum* interferes with monocyte-mediated inflammation, but the underlying molecular mechanisms remain unclear. Therefore, the aim of the current study was to investigate the transcriptional basis of the *A. suum*-suppressed monocyte phenotype using a systems immunology approach. Our data show comparable global gene expression profiles between *Ascaris* single- and coinfected pigs, both distinct from *Salmonella*-infected pigs. However, further analysis into co-expression profiles revealed downregulation of key genes involved in inflammatory signaling modules (AP-1 activation and MAPK pathways), majorly in the coinfected pigs relative to *Salmonella* single infected pigs. This suggests that while overall transcriptional responses to *Ascaris* infection are conserved, coinfection may be associated with additional suppression of proinflammatory transcriptional programs, highlighting disrupted transcriptional networks underlying helminth-mediated immune suppression.

## Materials and methods

### Animal experimental infection

The experimental study was implemented in accordance with the European Convention for the Protection of Vertebrate Animals used for Experimental and other Scientific Purposes and the German Animal Welfare Law and is reported in accordance with ARRIVE guidelines 2.0. The study was ethically approved by the Berlin State Office of Health and Social Affairs (approval number G0212/20).

Experimental infection, blood sampling and PBMC isolation was previously described ^24^. Briefly, the study consisted of four experimental groups of six pigs (hybrid German landrace and large white, aged 9 weeks at necropsy) each: controls (Ctrl), *A. suum* single-infection (As; 4 x 2000 infective eggs), *S.* Typhimurium single-infection (ST; 10^7^ CFU) and *A. suum- S.* Typhimurium coinfection (AsST) where pigs were first infected with *Ascaris* (4 x 2000 infective eggs), then subsequently infected with *Salmonella* (10^7^ CFU) 7 days later as described ^24^. Each group consisted of three males and three females. PBMCs were isolated from blood collected by heart puncture at the following dissection timepoints: 14 days post-infection (dpi) for *A. suum* and 7 dpi for *S.* Typhimurium ^24^.

### mRNA extraction and RNA sequencing

Frozen PBMCs were thawed at room temperature and washed in RPMI with 1% fetal calf serum (FCS), 100 U/ml penicillin and 100μg/mL streptomycin (PAN-Biotech). Thereafter, RNA was extracted from 5 x 10^6^ PBMCs using the innuPREP DNA/RNA Mini Kit (AJ Innuscreen GmbH, Berlin, Germany) and RNA concentration determined using Qubit™ RNA HS Assay Kit (ThermoFisher, USA). Poly-A enrichment, library preparation, and quality control were performed by Biomarker Technologies (BMK) GmbH (Münster, Germany) for mRNA sequencing on the Illumina NovaSeq 6000 platform (PE 150, 9 Gb/sample). The mRNA sequencing data contained 795.64 million reads with an average yield of 9.93 Gb per sample and at least 30 M reads per sample.

### Sequence pre-processing and alignment

Quality control of the sequences was done using fastQC (v0.12.1) while adapter removal and trimming of low-quality reads was done using fastp (v0.23.4) ^29^. The sequences were further aligned against the Ensembl’s porcine reference genome *Sus scrofa* (v11.1) using STAR aligner (v2.7.11b) ^30^. The read abundance levels were estimated at gene level using featureCounts (v2.0.6) ^31^. Further downstream analyses were carried out in R (v4.4.0) ^32^.

### Differential gene expression analysis

Quantified abundance estimates (count matrix) were quality controlled for counts less than 10. DEseq2^33^ (v1.48.1) was applied for normalization and differential gene expression using the incorporated geometric mean method and negative binomial distribution. Differentially expressed genes (DEGs) were determined by applying a threshold cutoff of |Log_2_foldchange| ≥ 1 & padj < 0.05 (p-value adjusted for multiple testing using Benjamini-Hochberg).

### Gene co-expression network analysis

DESeq2 variance-stabilized counts were filtered to the top 8,000 most variable genes for downstream co-expression network analysis using the WGCNA (v.1.73) ^34^ package. A soft-threshold power (*β* = 10) was chosen from scale-free topology fit. Gene similarity (Pearson correlation) was transformed into an adjacency matrix and clustered via signed TOM (min module size = 30, merge cut height = 0.25). Resulting co-expression clusters (“modules”) were color-labeled, with unassigned genes grouped into grey. For each module, a module eigengene (ME; the first principal component of the gene expression values) was calculated to summarize the dominant expression pattern. MEs were then correlated with metadata (external traits), including infection status and monocyte frequencies obtained from flow cytometry data (mean ± SD: uninfected controls, 3.2 ± 1.98; *A. suum* single-infected, 1.95 ± 0.55; coinfected, 1.55 ± 0.67; *S.* Typhimurium single-infected, 6.75 ± 2.19) [23]. Infection status, being categorical, was encoded separately for each infection type (e.g., *Ascaris* single-infected = 1, all others = 0), whereas monocyte frequencies, as continuous quantitative data, were included directly as one variable across all samples. This approach allowed us to test whether module expression patterns tracked with monocyte abundance without splitting monocytes by infection status. Group-level comparisons of ME values were then performed to visualize how module expression trends paralleled observed monocyte frequency changes across infection groups, thereby integrating immune cell abundance with transcriptional modules in a signed weighted network framework.

### Functional analysis

Functional analyses were conducted using the clusterProfiler package (v4.16.0) ^35^. Gene set enrichment analysis (GSEA) was applied on significant contrasts and key co-expression modules using the gene ontology (GO) and Kyoto Encyclopedia of Genes and Genomes (KEGG) databases. In addition, over-representation analysis was applied on the DEGs. Protein-protein interaction network analysis was conducted using the STRINGdb package v.2.20 ^36^, configured for STRING version 12.

## Results

### Opposing transcriptional profiles in PBMCs of single infections with *Ascaris* versus *Salmonella*

To explore the systemic immune imprint of each pathogen, we first compared PBMC gene expression profiles of single-infected pigs to uninfected controls. Only one gene (*ENSSSCG00000055451)* was differentially expressed in *A. suum* single-infected pigs (14 days post infection) when compared to uninfected controls (**Fig. 1A**), an observation consistent with our previously reported weak systemic response and strong local migratory-dependent immune response in an experimental acute *A. suum* infection in juvenile pigs ^23^. Similarly, *S.* Typhimurium single-infected pigs (7 days post infection) showed no significant transcriptional changes in PBMCs compared to uninfected controls (**Fig. 1A**). This finding suggested that neither infection alone elicited strong systemic transcriptional shifts in circulating immune cells. As the PBMCs were analyzed *ex vivo* without additional *in vitro* restimulation, the data capture only the baseline systemic imprint of infection, and subtle differences might have become apparent upon restimulation.

**Fig. 1:**
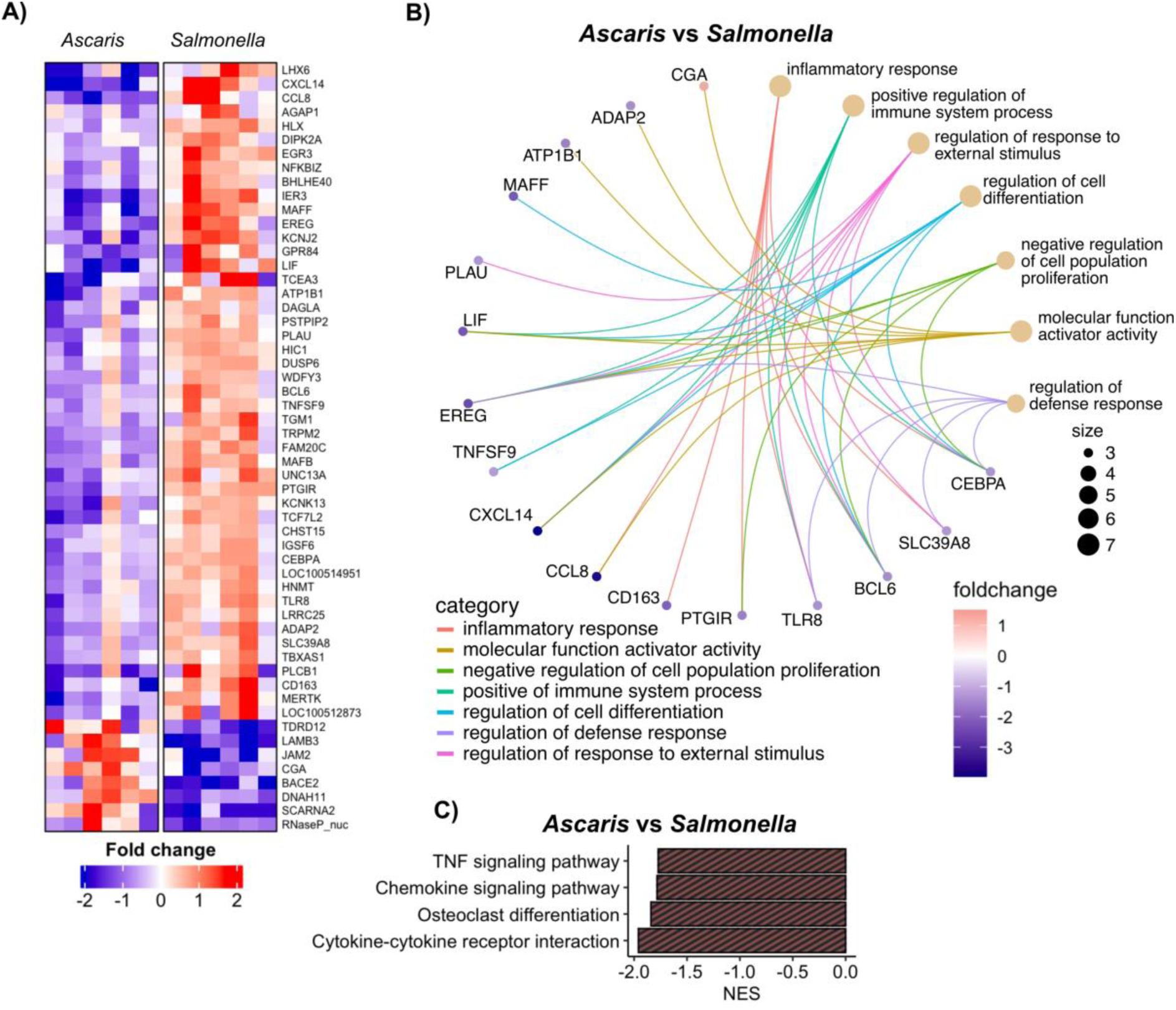
Differential gene expression analysis between PBMCs of *Ascaris* single-infected and *Salmonella* single-infected pigs. **A)** Venn diagram showing the DEGs in unstimulated PBMCs of *A. suum* (As)-infected, *S.* Typhimurium (ST)-infected compared to the uninfected control (ctrl) group (As vs ctrl; ST vs ctrl) and compared to each other (As vs ST). **B)** Heatmap illustrating 55 DEGs in PBMCs of *Ascaris* single-infected pigs compared to *Salmonella* single-infected pigs. Red indicates the positive log_2_fold change (upregulation) whilst blue indicates negative log_2_fold change (downregulation). x axis; sample identification, y axis; genes identified. **C)** C-net plot of top over-represented GO terms indicated by the brown dots and the corresponding DEGs within the terms. The log fold changes of the DEGs are indicated by the red-blue gradient with increasing intensity corresponding to increase (red) or decrease (blue) in foldchange. **D)** Bar plot of significantly enriched KEGG pathways after GSEA analysis. A negative normalized enrichment score (NES) indicates suppression of pathways in PBMCs of *Ascaris* single-infected pigs compared to the *Salmonella* single-infected pigs. The y axis indicates the KEGG pathways, and the x axis indicates the normalized enrichment score. ST: *S.* Typhimurium, As: *A. suum*, ctrl: Uninfected control.

A comparison between the two single-infected groups revealed a clearer transcriptional distinction. Pigs infected for 14 days with *A. suum* exhibited 54 downregulated and 14 upregulated genes relative to *S.* Typhimurium-infected counterparts (**Fig. 1B; Fig. S1A**). Among the downregulated genes, *CXCL14, TLR8, SLC39A8, CD163, CCL8* and *TNFSF9* are associated with monocyte activation and chemotaxis ^37–40^. Functional enrichment analyses further supported this anti-inflammatory bias. Downregulated genes in *A. suum*-infected pigs were enriched in processes related to inflammatory responses, defense regulation, and cytokine signaling (**Fig. 1C, Fig. S1B)**. Complementary KEGG pathway analysis via GSEA revealed suppression of TNF signaling, chemokine signaling, osteoclast differentiation, and cytokine-cytokine receptor interactions in *A. suum* PBMCs (**Fig. 1D)**.

Taken together, these findings suggest that while baseline PBMC transcriptional profiles in single infections were largely indistinguishable from uninfected controls, subtle yet biologically meaningful differences emerged when comparing the two single infections. Specifically, *A. suum* infection induced a systemically muted, anti-inflammatory profile characterized by the downregulation of monocyte-related genes, in contrast to the inflammatory signature associated with *S.* Typhimurium. This likely reflects the localized nature of both infections and indicates that comparative analysis, rather than control-based contrasts alone, may be more revealing of early immune modulation.

### *Ascaris* coinfection dampens systemic inflammatory responses to *Salmonella*

Building on our observation of minimal transcriptional changes in single-infected pigs relative to controls, we next investigated how *A. suum* and *S.* Typhimurium coinfection shaped systemic immune responses in PBMCs. We began by comparing PMBCs from coinfected pigs to the uninfected group. Only seven genes (*ENSSSCG00000006932, ENSSSCG00000055451*, *SHISA2*, *ENSSSCG00000054605, OSM*, *ENSSSCG00000050752, ZBTB16)* were differentially expressed, highlighting a limited systemic transcriptional response to coinfection; similarly observed in the single infections (**Fig. 2A**). Moreover, coinfected pigs showed no significant differences in PBMC gene expression compared to *A. suum* single-infected pigs (**Fig. 2A**), reinforcing the similarity in their systemic immune signatures. In contrast, 126 genes (20 upregulated and 106 downregulated) were differentially expressed in coinfected pigs in comparison to *S.* Typhimurium single-infected pigs (**Fig. 2B; Fig. S1C**).

**Fig. 2:**
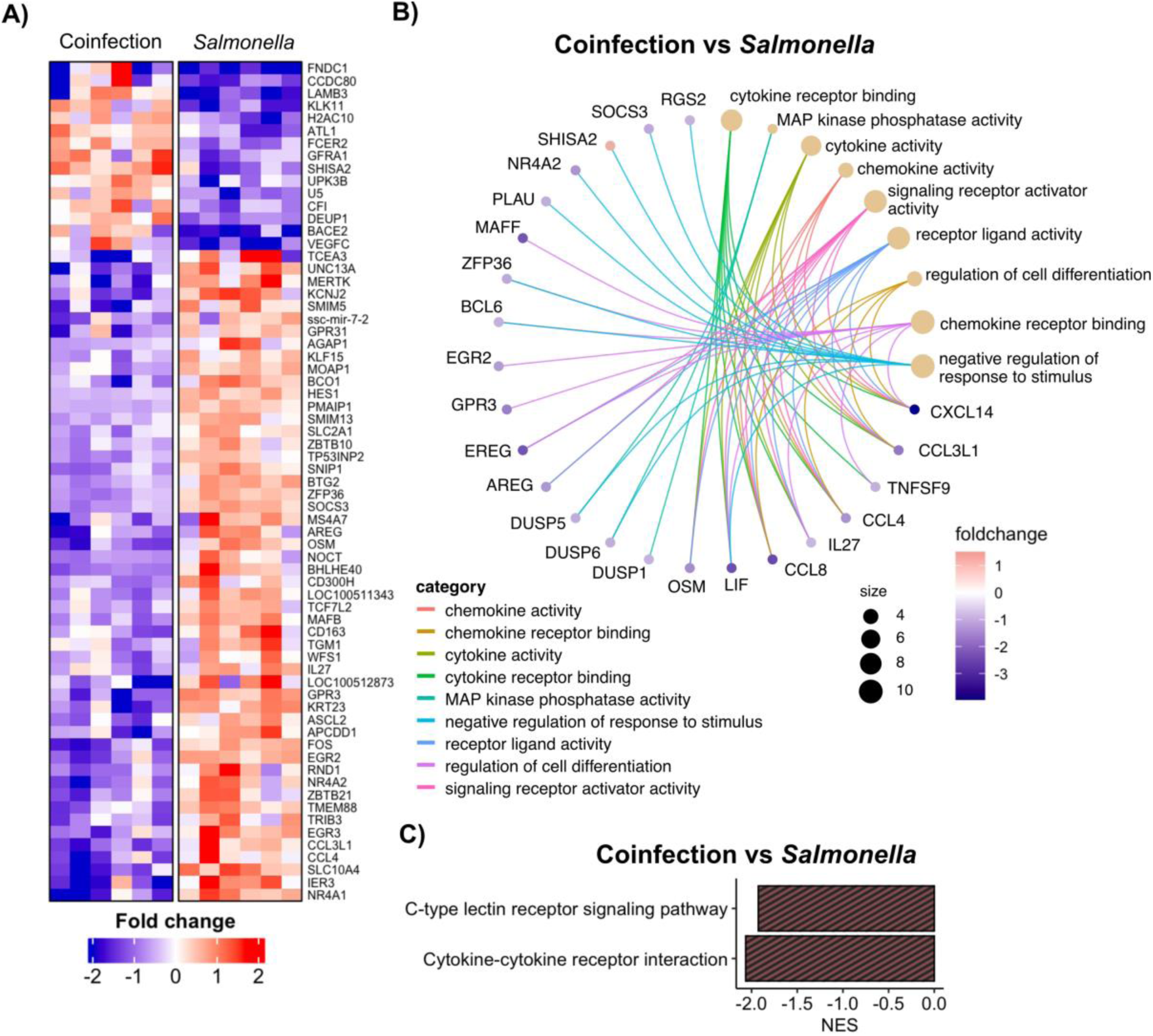
Genes involved in cytokine and chemokine activity are suppressed in coinfected pigs in comparison to *Salmonella* single-infected pigs. **A)** Venn diagram showing the DEGs in coinfected (AsST) compared to the uninfected control (ctrl), *S.* Typhimurium (ST)-infected or *A. suum* (As)-infected (AsST vs ctrl; AsST vs ST; AsST vs As). **B)** Heatmap illustrating 67 DEGs in PBMCs of coinfected pigs compared to *Salmonella* single-infected pigs. Red indicates positive log_2_fold change (upregulation) whilst blue indicates negative log_2_fold change (downregulation). x-axis; sample identification, y-axis; genes identified. **C)** C-net plot of top over-represented GO terms indicated by the brown dots and the corresponding DEGs within the terms. The log fold changes of the DEGs in cnet plots are indicated by the blue-red gradient with increasing intensity corresponding to increase (red) or decrease (blue) in foldchange. **D)** Bar plot of significantly enriched KEGG pathways after GSEA analysis. A negative normalized enrichment score (NES) indicates suppression of the pathways in the coinfected pigs compared to *Salmonella* single-infected pigs. ST: *S.* Typhimurium, As: *A. suum*, ctrl: Uninfected control.

Functional enrichment of these 126 DEGs revealed suppression of immune-related processes and signaling functions in coinfected pigs. Downregulated genes were enriched in GO terms for biological processes such as ‘negative regulation of stimulus response’ and ‘regulation of cell differentiation’, and molecular functions including ‘cytokine receptor binding’ and ‘signaling receptor activator activity’ (**Fig. 2C; Fig. S1D)**. Markedly, cytokine-cytokine receptor interaction pathways were suppressed in coinfected pigs, mirroring the pattern observed in *A. suum* single-infected animals (**Fig. 1D**; **Fig. 2D**), implicating a helminth-driven dampening of cytokine-mediated communication networks crucial for coordinating antibacterial responses. Additionally, we observed downregulation of the C-type lectin receptor (CLR) signaling pathway **(Fig. 2D**), a pathway known to be critical for antimicrobial defense ^41^. GO terms enriched in *S.* Typhimurium single-infected pigs, such as MAPK cascade activation, immune response, and responses to biotic stimulus, were notably absent in the coinfected group (**Fig. S1D)**, further pointing to a repressed inflammatory program.

Collectively, these results reveal that the PBMC transcriptional profile of coinfected pigs aligns more closely with that of *A. suum*-infected pigs than with *S. Typhimurium* single-infected counterparts. This convergence suggests that a previous and active *A. suum* infection dominantly shaped the systemic immune landscape in coinfected pigs, suppressing subsequent *Salmonella*-driven transcriptional programs, including key cytokine and innate immune signaling pathways, thereby potentially compromising antibacterial immunity.

### Monocyte-associated gene modules reveal inflammatory networks disrupted by coinfection

Comparative analysis revealed no DEGs in either *S.* Typhimurium or *A. suum* single-infected pigs when compared to uninfected controls (**Fig. 1A**). However, a comparison between the two single infections revealed that *A. suum* infection was associated with suppression of TNF signaling and reduced expression of monocyte-associated chemokines relative to *S. Typhimurium* infection. These subtle yet biologically meaningful differences pointed to infection-specific immune modulation that may not be captured by standard DEG thresholds. To gain a broader systems-level view of host responses, we applied signed weighted gene co-expression network analysis (WGCNA) ^42, 43^, clustering genes into modules based on their shared expression patterns and summarizing each with an ME. Correlating these MEs with infection status and monocyte frequencies obtained from flow cytometric data enabled us to assess how transcriptional modules tracked with immune cell abundance across different infection states.

Co-expression network analysis identified 21 gene co-expression modules, whose MEs were correlated with infection status and monocyte frequencies (**Fig. 3A**). Significant module-trait associations were defined as those with |r| > 0.5 and p-value < 0.001. Strikingly, the *Salmonella* single-infected group showed a strong positive correlation with the blue module (*r*=0.59, *p*=0.002), which also correlated with monocyte frequencies (*r*=0.79, *p*=4e-06). Monocyte frequencies were additionally associated with the brown (*r*=0.66, *p*=5e-04) and lightcyan (*r*=0.63, *p*=9e-04) modules **(Fig. 3A),** suggesting co-regulated networks underlying innate immune activation. In contrast, the coinfection group showed a distinct transcriptional profile, aligning most closely with the red module (*r*=0.54, *p*=0.006). Neither the uninfected nor *Ascaris* single-infected groups showed significant module associations, reinforcing the impression of transcriptional dampening or lack of targeted immune activation in these groups. (**Fig. 3A**). Analysis of the ME values in the blue module, across infection groups, showed a marked upregulation in the *Salmonella* single-infected group relative to both *Ascaris* single- and coinfected groups **(Fig. 3B).** This pattern mirrored monocyte frequencies, which increased during Salmonella infection but declined in the other two groups. Meanwhile, the red module, dominant in coinfection, was markedly repressed in the *Salmonella* group **(Fig. 3C)**. Together, these patterns highlight distinct transcriptional landscapes shaped by single versus coinfection, with monocyte-driven responses featuring prominently during *Salmonella* infection.

**Fig. 3:**
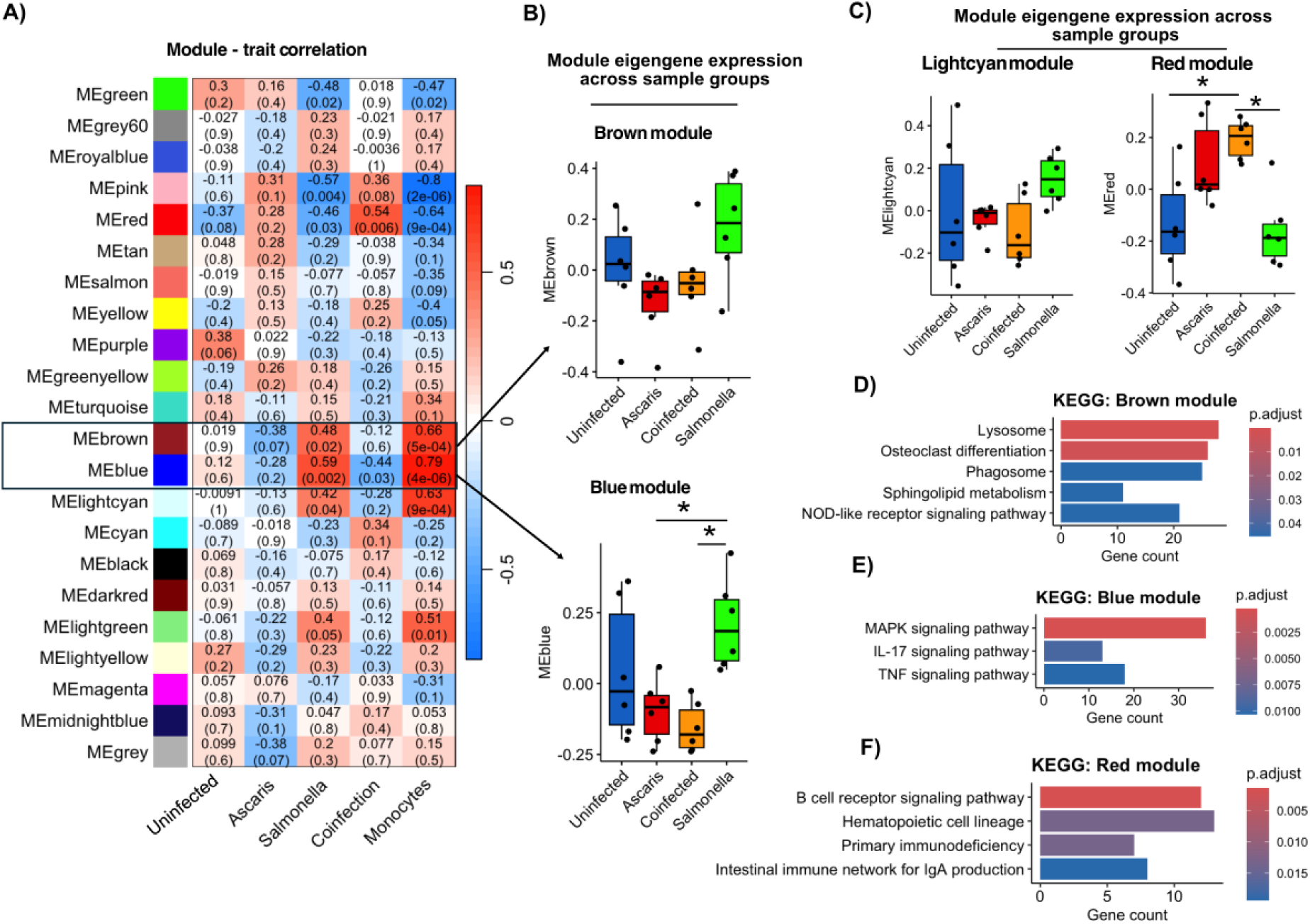
Inflammatory and phagosome/lysosome modules associated with anti-*Salmonella* monocytic response. **A)** Weighted gene-co-expression network analysis (WGCNA) module–trait heat-map. Each cell shows the Pearson correlation (color intensity) and corresponding p-value (in brackets) between module eigengenes (ME; rows) and clinical traits (columns). Red indicates positive correlation while blue indicates negative correlation. Boxplots **(B, C)** showing ME values of brown, blue, lightcyan and red modules across uninfected (blue), *Ascaris* infected (red), coinfected (orange) and *Salmonella* infected (green) pigs. The y-axis; ME value and the x-axis; infection groups. Bar plots of significantly enriched KEGG pathways of the brown **(D)**, blue **(E)** and red **(F)** modules. The y axis of the bar plots represents the KEGG pathways whilst the x axis represents the number of genes within the pathway. The gradient color of the bars indicates significance with red as most significant.

To explore biological functions of these modules, we performed KEGG enrichment analysis. The brown module, which correlated with monocyte frequencies, was enriched for genes involved in osteoclast differentiation and lysosome/phagosome pathways, hallmarks of myeloid cell activation (**Fig. 3D).** The blue module, which is also monocyte-associated and dominant in the *Salmonella* group, was enriched for proinflammatory signaling cascades (TNF, IL-17, and MAPK; **Fig 3E)**. In contrast, the red module, upregulated in coinfection, showed enrichment for B cell receptor signaling and mucosal networks such as Immunoglobulin (Ig)A production (**Fig. 3F)**, indicating adaptive/mucosal responses. The light cyan module, though monocyte-associated, lacked significant functional enrichment, suggesting limited functional cohesion.

Together, these results highlight how distinct infections modulate transcriptional programs differently, *Salmonella* driving inflammatory monocyte signatures, coinfection engaging mucosal immunity, and *Ascaris* exerting a more silent or suppressive effect.

### Driver genes uncover suppressed MAPK/AP-1 inflammatory regulators during coinfection

The blue and brown modules, enriched for monocyte functions and inflammatory pathways (MAPK, TNF, IL-17) alongside phagosome/lysosome activity, represent key innate immune networks and were selected for further investigation. To better understand the molecular drivers within these networks, we focused on genes strongly correlated to monocyte frequencies. By correlating gene significance (GS) with module membership (MM), we found robust relationships in both modules (brown: *r*=0.76, *p*=9.8e-163 (**Fig. S2A);** and blue: *r*=0.79, *p*=1e-200 (**Fig. 4A**), suggesting that genes central to these modules mirrored shifts in monocyte frequencies across infection groups. Applying stringent thresholds of |GS| ≥ 0.7 and MM ≥ 0.7, we identified 47 driver genes in the brown module and 102 in the blue.

**Fig. 4:**
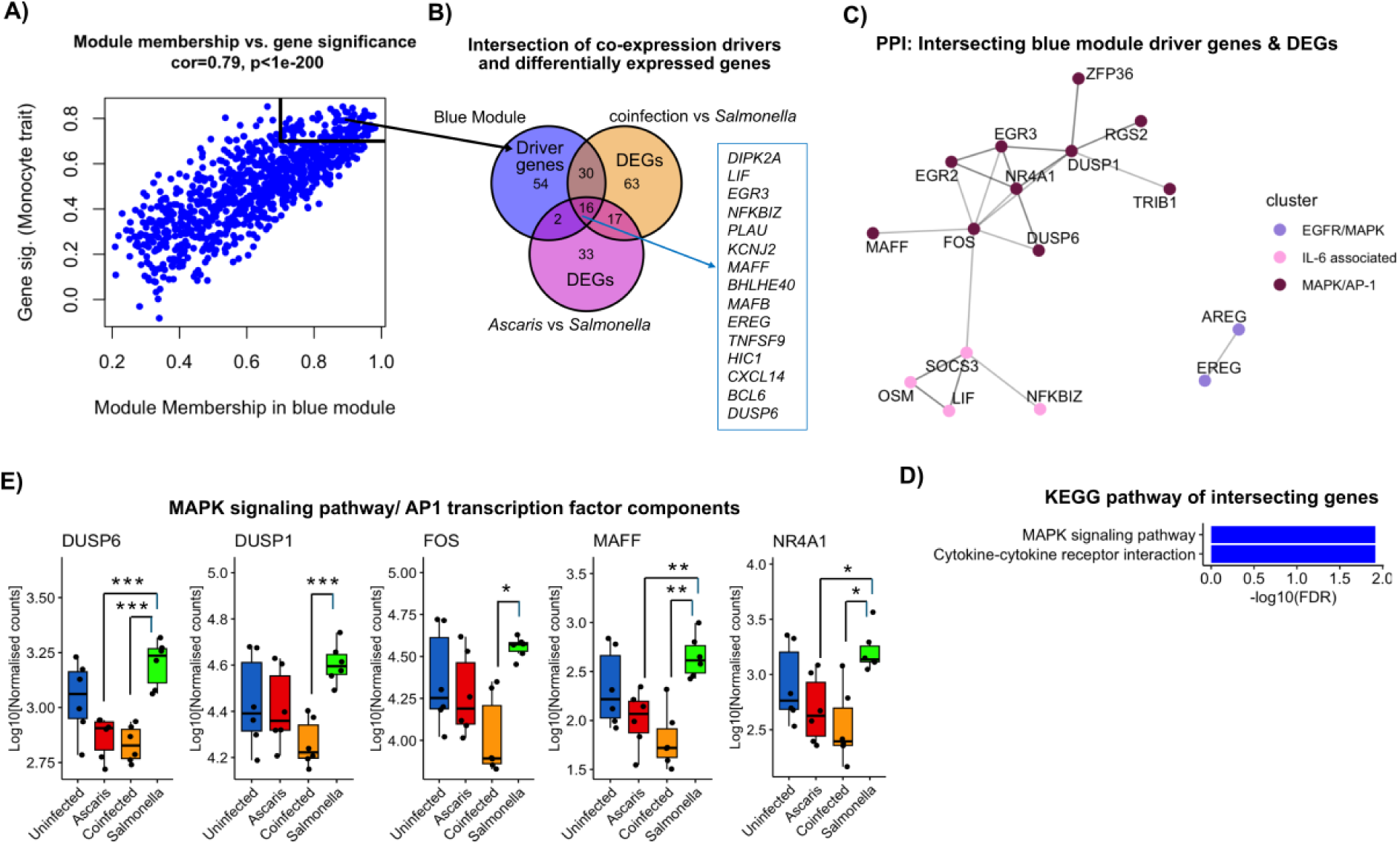
Integration of co-expression analysis, differential expression, and functional enrichment identifies MAPK signaling as a key pathway. **A)** Scatter plots depicting the correlation between module membership (x axis) and gene significance (y axis) for the blue module (monocyte trait-associated), with a strong correlation (0.79, *p* < 1e-200). **(B)** Venn diagram showing intersecting genes between blue module driver genes and DEGs from *Ascaris* vs *Salmonella* and coinfected vs *Salmonella* contrasts. **(C)** STRING protein–protein interaction network of intersecting genes, revealing a tightly connected subnetwork. Clusters are depicted by the green, red and blue nodes (bubbles). K-means clustering was used. The edges (lines) are colored according to interaction. Dotted lines depict connected clusters. Black line denotes co-expression evidence; magenta denotes experimental evidence; green denotes neighborhood evidence; cyan blue denotes database evidence. **(D)** Bar plots showing KEGG pathway enrichment analysis of intersecting genes in the blue module. **(E)** Boxplots display log₂-normalized expression of selected MAPK/AP1-related genes across uninfected (blue), *Ascaris* infected (red), coinfected (orange) and *Salmonella* infected (green) pigs.

To explore which of the drivers were different between infections, we overlapped these driver genes with DEGs from *Ascaris* vs *Salmonella* and coinfection vs *Salmonella*. In the brown module, only nine of 47 driver genes overlapped with DEGs from *Ascaris* vs *Salmonella* and four intersected with DEGs from the coinfection vs *Salmonella* respectively (**Fig. S2B**). Of these, three shared genes (*TGM1*, C*D163* and *HLX)* were consistently upregulated in *S.* Typhimurium single-infected pigs. While indicative of monocyte activation, their sparse overlap and weak PPI connectivity suggested a limited network footprint.

In the blue module, a more substantial set of driver genes overlapped with DEGs. Eighteen driver genes were shared with DEGs distinguishing *Ascaris* from *Salmonella* infection, while 46 overlapped with DEGs from the coinfection vs *Salmonella* comparison, highlighting a broader disruption of monocyte-associated transcriptional programs during coinfection (**Fig. 4B**). This larger gene set allowed for a more coherent picture to emerge. PPI analysis of the 46 genes revealed three interconnected subnetworks **(Fig. 4C)**. At the center was a cluster of transcriptional regulators, including *FOS, EGR2, EGR3,* and *MAFF,* alongside phosphatases *DUSP1* and *DUSP6*. These nodes were linked through *FOS* to a secondary network rich in IL-6 family signaling, *LIF, OSM, SOCS3,* and *NFKBIZ*, all known mediators of inflammation and immune modulation. A third, distinct cluster comprised the epidermal growth factor receptor (EGFR) ligands *AREG* and *EREG*.

Interestingly, the PPI analysis highlighted a densely connected cluster involving AP-1-associated transcription factors (*FOS*, *NR4A1*, *MAFF*, *MAFB*) and MAPK related genes (*FOS*, *DUSP1*, *DUSP6*, *AREG*, *EREG*). KEGG pathway enrichment of these intersecting genes confirmed MAPK signaling as a significantly enriched pathway (adjusted *FDR* < 0.0121), alongside cytokine–cytokine receptor interaction (**Fig. 4D**). Direct comparison of these genes across the groups revealed reduced levels of AP-1 components (*FOS, MAFF, NR4A1*) as well as the MAPK regulators *DUSP1* and *DUSP6* in coinfected pigs in comparison to *Salmonella* single infected pigs (**Fig. 4E**). Additional cytokine-related genes, including *CCL3L1*, *CXCL14*, *SOCS3* and *OSM* also showed infection-specific suppression (**Fig. S2C)**. These findings suggest that *A. suum* actively antagonizes the proinflammatory transcriptional program otherwise induced by *S. Typhimurium*, reshaping the immune landscape at the network level.

Together, these analyses highlight the blue module as a central hub of monocyte-driven immune responses to *S. Typhimurium*, characterized by coordinated expression of inflammatory transcription factors, signaling molecules, and cytokine regulators. The marked downregulation of key AP-1 and MAPK-associated genes during coinfection points to a suppressed inflammatory program, likely contributing to the attenuated monocyte activation observed in coinfected pigs. These findings suggest that coinfection with *A. suum* not only dampens transcriptional responses but may also disrupt critical regulatory circuits required for effective anti-*Salmonella* monocyte immunity.

## Discussion

Our study provides mechanistic insight into how *A. suum* infection compromises anti-*Salmonella* immunity by modulating monocyte transcriptional responses. Building on our earlier observation of impaired monocyte activity and increased *Salmonella* burden in coinfected pigs ^24^, we now show that *A. suum* suppresses AP-1/MAPK-driven inflammatory networks in PBMCs of coinfected pigs. While *S. Typhimurium* infection alone triggered robust monocyte-associated responses, including activation of TNF, IL-17, and MAPK signaling, a previous and ongoing *A. suum*-infection disrupted these pathways, leading to a muted inflammatory transcriptional program. These findings shed light on the molecular circuitry underlying helminth-mediated immune suppression and offer a new framework for understanding the impact of coinfections on bacterial clearance.

Anti-*Salmonella* proinflammatory cytokine responses are driven by transcription factors such as AP-1 and NF-κB ^44^. AP-1, typically formed as a *FOS–JUN* heterodimer, orchestrates immune cell activation, proliferation and differentiation ^45^ and is enhanced by MAPK cascades ^46^, which are triggered by microbial infections, cytokines or growth factors such as amphiregulin (AREG) and epiregulin (EREG). IL-4 pretreatment has been shown to inhibit *FOS* expression in LPS-stimulated monocytes and macrophages ^47, 48^, consistent with IL-4-driven polarization toward an M2 phenotype that favors intracellular *S.* Typhimurium replication ^21, 22, 49^. In line with this, we observed significantly elevated frequencies of CD4^+^IL-4^+^ T cells in the PBMCs of *A. suum* infected pigs, suggesting a host environment biased toward IL-4-mediated immune regulation ^24^. At the transcriptional level, coinfected pigs displayed downregulation of AP-1-associated transcription factors (*FOS*, *MAFF*, *NR4A1*), MAPK regulators (DUSP1/6) and EGFR ligands (*AREG/EREG*) ^46, 50, 51^, suggesting a MAPK environment in which both activating inputs and feedback control are impaired. We showed earlier that helminths are also capable of exploiting the MAPK pathway, inducing *DUSP1* in monocytes and macrophages thereby reinforcing an IL-10 mediated immunosuppressive state ^52^. In our setting here, single *A. suum* infection did not modulate expression levels of DUSP1, whereas single *S.* Typhimurium infection strongly induced it, reflecting feedback control of inflammation. Strikingly, this induction was lost under coinfection, coinciding with the failure of monocytes to produce TNF-α after IL-12 stimulation ^24^.

The MAPK and AP-1 signaling pathways were further linked to the IL-6 cytokine signaling family genes (*LIF* and *OSM*) via *FOS* and the negative regulator *SOCS3* gene. Markedly, *SOCS3* transcription is directly influenced by AP-1 components, with its promoter harboring binding sites for both *FOS* and *JUN* ^53^, highlighting the integration of AP-1 activity in cytokine signaling control. In this study, PBMCs from coinfected pigs exhibited significantly lower expression of *LIF, OSM* and *SOCS3* genes relative to PBMCs from *S.* Typhimurium single-infected pigs. *SOCS3* gene is strongly induced by bacterial LPS ^54, 55^ and is a known promoter of M1 macrophage polarization, whereas a higher *SOCS1/SOCS3* gene ratio has been associated with an M2-like, anti-inflammatory macrophage phenotype ^54, 56, 57^. In line with this, Liu *et al.* demonstrated that *SOCS3*-deficient macrophages failed to mount M1 responses despite LPS stimulation ^54^. These findings highlight the functional relevance of *SOCS* gene expression in fine-tuning cytokine responses through regulation of STAT activation in the context of gp130-mediated signals ^58^.

The IL-6 family cytokines *OSM* and *LIF* (downregulated in PBMCs of coinfected pigs) share glycoprotein130 receptor signaling with IL-6 ^59^, thus we postulated non-competitive binding of IL-6 to the gp130 receptor. Reduced *SOCS3* expression likely disrupted negative feedback on IL-6/gp130/STAT3 signaling [60], potentially exacerbating IL-6-mediated immunosuppression while impairing LPS-induced TNF-α production [61]. Accordingly, we previously found decreased TNF-α response in restimulated PBMCs of coinfected pigs ^24^. Furthermore, research by Pagin *et al* demonstrated that inhibition of AP-1 activity, through reduced *FOS* expression, resulted in lower *SOCS3* expression ^53^. This suggests that diminished *FOS* expression in coinfected pigs likely disrupted AP-1 complex formation, thereby reducing *SOCS3* expression. The resultant decline in *SOCS3*-mediated regulatory activity likely contributed to an impaired anti-*Salmonella* inflammatory response in the coinfected pigs.

Consistent with our previous findings of diminished monocyte responsiveness in PBMCs from coinfected pigs ^24^, we observed reduced expression of chemokine genes critical for monocyte recruitment (*CCL8* and *CXCL14*) and macrophage-mediated antimicrobial activity (*CCL3L1*) in coinfected pigs relative to those infected solely with *S.* Typhimurium*. CXCL14* expression is largely restricted to monocytes and B cells ^60^ and has been implicated in the regulation of AP-1 and NF-κB signaling pathways, as well as direct antimicrobial activity ^39^. *CCL3L1* encodes macrophage inflammatory protein (MIP)-1α, a proinflammatory chemokine primarily secreted by monocytes and macrophages and known to be suppressed by Th2 cytokines such as IL-4, IL-10 and IL-13 ^61, 62^. These findings further support a Th2-skewed, anti-inflammatory environment in the coinfected pigs. In addition, *TNFSF9*, which encodes CD137 Ligand signaling, a key costimulatory molecule involved in monocyte activation, dendritic cell differentiation, and proinflammatory cytokine induction along with proinflammatory cytokine induction ^63^, was significantly downregulated in both *A. suum* single- and coinfected pigs relative to those infected with *S.* Typhimurium only. The concerted suppression of these monocyte-attracting and activating signals likely contributes to the impaired peripheral monocyte inflammatory landscape observed in coinfection.

Our findings further support the notion of dampened inflammatory responses in *Ascaris* single- and coinfected pigs, evidenced by the low expression of *HLX* and *CD163*. The transcription factor *HLX* promotes Th1 differentiation by inducing IFN-γ in immature T helper cells and cooperating with T-bet ^64, 65^, while *CD163,* a hemoglobin scavenger receptor, is positively associated with inflammatory cytokine production including IL-6 and IL-10 ^66^. Suppression of these genes suggests impaired Th1 skewing and altered macrophage activation states in helminth-exposed pigs.

We acknowledge several limitations in this study. First, the relatively low infection dose of *S.* Typhimurium may limit the extent of detectable systemic immune responses. Specifically, the *S.* Typhimurium dose used (10⁷ CFU) resulted in an asymptomatic, subclinical infection ^24^, in contrast to studies using >10⁹ CFU that induce overt clinical symptoms of fever and diarrhea by 2 dpi ^67^. However, subclinical carriage more closely mirrors natural infections in pigs, where both helminth and bacterial infections can persist without overt disease. Second, our transcriptomic analysis identified key immunomodulatory genes and pathways altered in PBMCs during *A. suum–Salmonella* coinfection, however these findings are based on computational inference. Functional studies are needed to confirm the roles of key genes such as DUSP1/6, *FOS*, *SOCS3*, and *CD163* in shaping monocyte responses during coinfection.

Collectively, our findings demonstrate that *A. suum* coinfection suppresses AP-1/MAPK-dependent transcriptional signatures in peripheral monocytes, thereby compromising anti-*Salmonella* inflammatory responses. More broadly, this work provides mechanistic insight into how helminth-induced immune modulation can impair host defenses against secondary bacterial pathogens.

## Data availability

Sequence data generated from this project will be made available at the NCBI Sequence Read Archive (SRA).

## Acknowledgments

This work was supported by German Research Foundation (Deutsche Forschungsgemeinschaft: DFG) grant HA 2542/11-2 and DFG GRK 2046 to SH. ZDM thanks Deutscher Akademischer Austauschdienst (DAAD) for the doctoral financial support [PIN: 91771723]. The authors would like to thank Beate Anders, Yvonne Weber, Bettina Sonnenburg, Marion Müller, and Christiane Palissa for methodological and technical support.

## Author contributions

SH, AM and ZDM conceived the study. ZDM, RH, RMM, LO, J. S-B and AM performed the experiments. ZDM curated, analyzed and visualized the data. ZDM and AM interpreted the data. SH and AM supervised the work. SH provided the funding and resources. ZDM wrote the original manuscript draft. ZDM, RH, RMM, LO, SR, JS-B, SH and AM reviewed and edited the manuscript draft. All authors have read and agreed to the published version of the manuscript.

