## Supplementary Figures for "Transcriptome of Peripheral Blood Mononuclear Cells Reveals Suppressed MAPK/AP-1 Pathways During *Ascaris-Salmonella* Coinfection in Pigs"

[#] Current Affiliation

[*] Corresponding author

 &

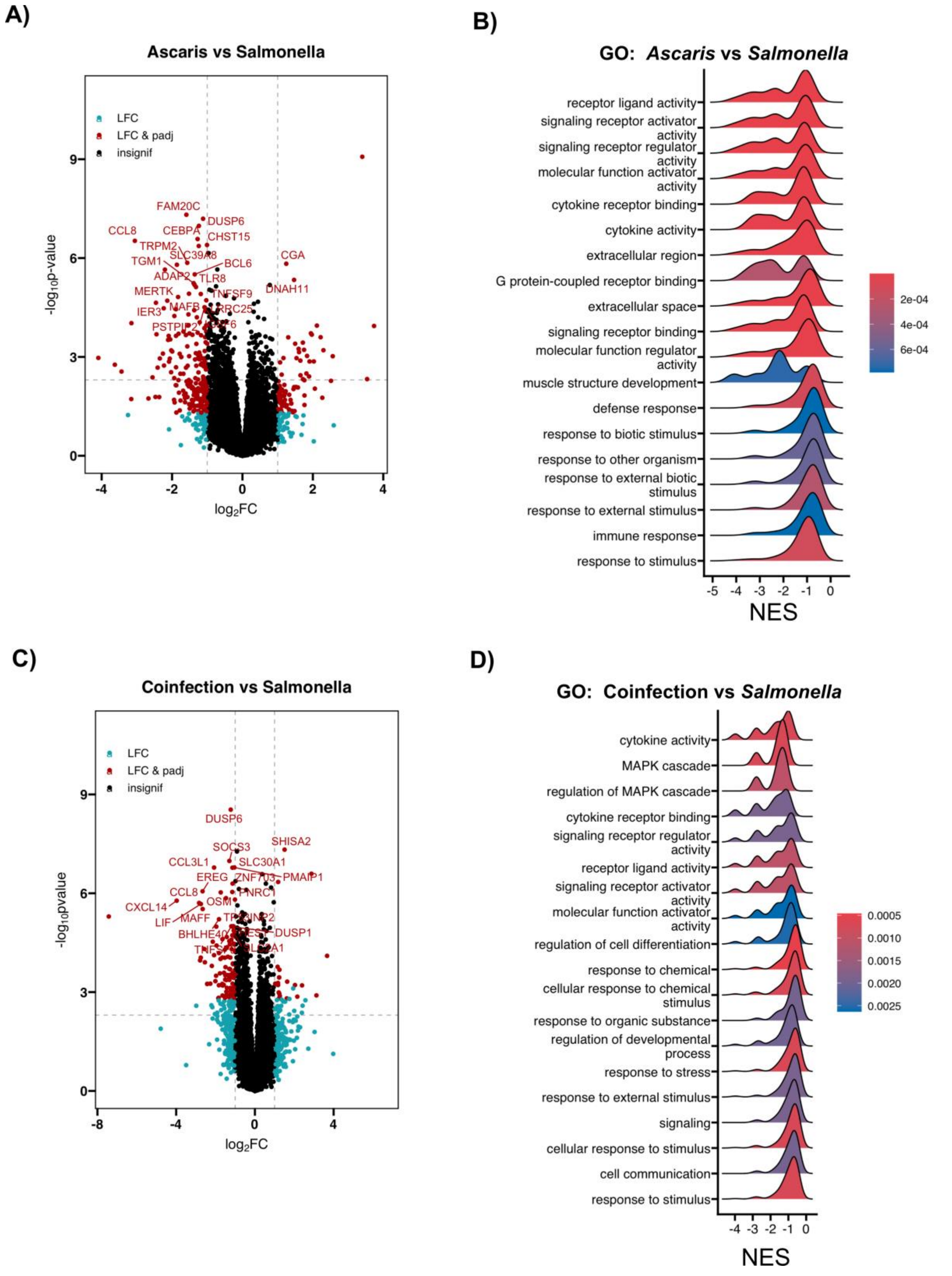


**Fig. S1**. **A)** Volcano plots depicting *Ascaris* vs *Salmonella* differentially expressed genes. The x-axis represents the $\log_2$fold change while the y-axis represents the $-\log_{10}$p-value. The top 20 DEGs are labelled. Black dots indicate insignificant genes; blue dots indicate genes at $|L_2FC| \geq 1$ whilst red dots indicate genes at padj < 0.05 & $|L_2FC| \geq 1$. **B)** Ridge plots showing gene ontology (GO) terms that significantly differ between PBMC transcriptomes of *Ascaris* infected and *Salmonella* infected pigs from GSEA analysis. The normalized enrichment score (NES) indicates whether a process is increased (positive NES) or suppressed (negative NES). The color gradient indicates the Benjamini-Hochberg adjusted p-value. **C)** Volcano plots depicting DEGs of coinfection vs *Salmonella* contrast. **D)** Ridge plots showing GO terms that significantly differ between PBMC transcriptomes of Salmonella single- and coinfected pigs from GSEA analysis.

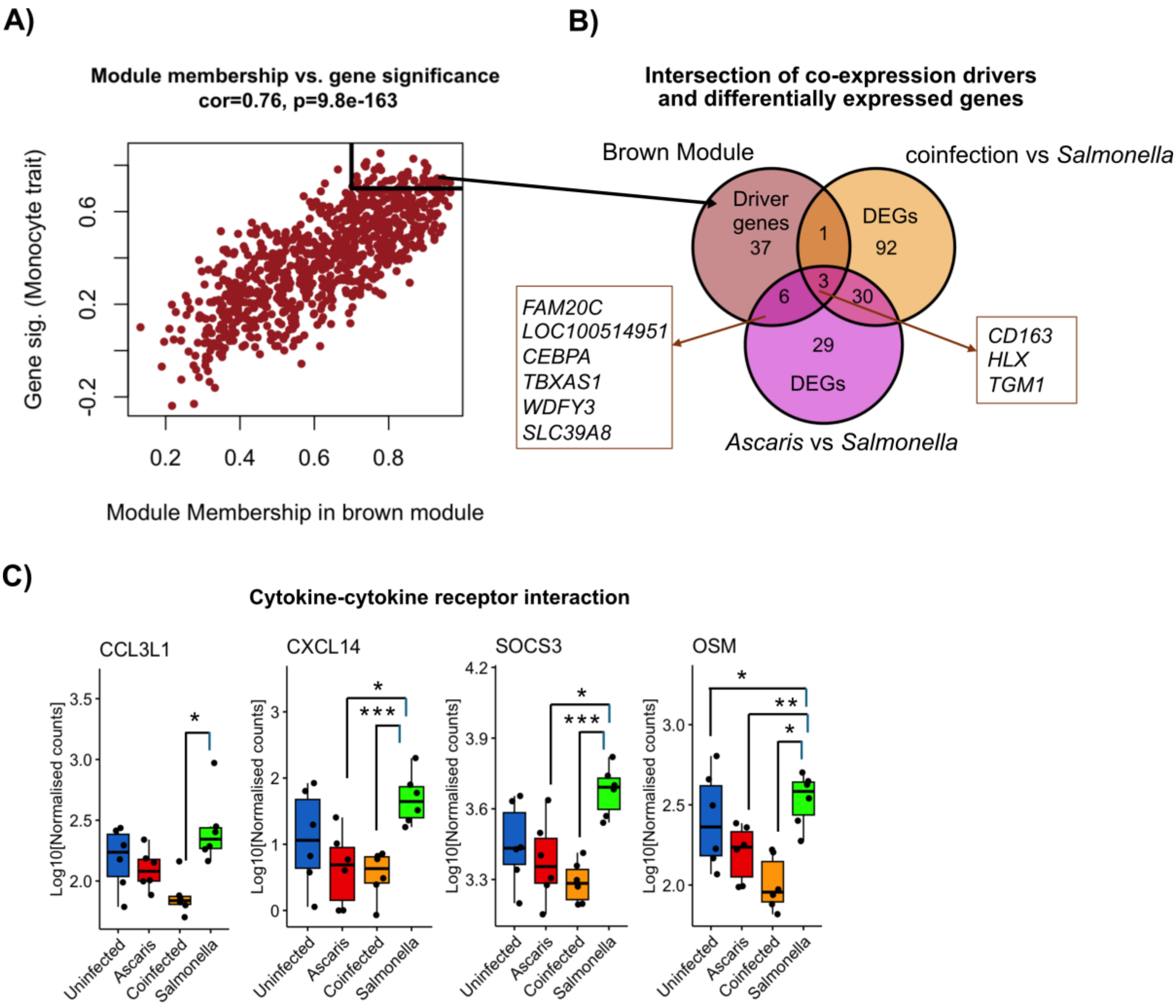


**Fig. S2. A)** Scatter plots depicting the correlation between module membership (x axis) and gene significance (y axis) for the brown module (monocyte trait-associated), with a strong correlation (0.76, $p$ =9.8e-163). **(B)** Venn diagram showing intersecting genes between brown module driver genes and DEGs from *Ascaris* vs *Salmonella* and coinfected vs *Salmonella* contrasts. **C)** Expression of cytokine signaling genes enriched in the cytokine–cytokine receptor interaction KEGG pathway. Boxplots display $\log_2$-normalized expression values of *CCL3L1*, *CXCL14*, *SOCS3* and *OSM* across infection groups.
